# Disturbance increases soil microbiome functional redundancy but decreases capacity for insurance via winnowed environmental responsiveness

**DOI:** 10.1101/2025.09.19.677300

**Authors:** Samuel E. Barnett, Ashley Shade

## Abstract

Redundancy is the capacity of coexisting populations to perform similar functions, supporting stable outputs under disturbance. The insurance hypothesis proposes that diverse communities are more resilient because they can respond to varied environmental stressors. Both redundancy and insurance likely shape resilience, but their relative contributions and potential interactions remain unclear. Here, we examined functional redundancy and potential for insurance in soil microbial communities disturbed by an ongoing underground fire in Centralia, Pennsylvania, USA, using seven years of deep metagenome sequencing from heated and reference sites. Analyses of taxonomic and functional diversity showed that functional redundancy increased with disturbance intensity, driven by declines in both diversity and average genome size. Biogeochemistry-relevant metabolisms, such as carbohydrate and nitrogen pathways, showed greater per-genome investment with disturbance intensity, indicating enhanced redundancy. However, transcription factor evenness – linked to community environmental responsiveness- was lower in heated soils. Thus, prolonged disturbance that imposes a strong selection can increase redundancy and near-term stability while simultaneously diminishing insurance potential against future stress, revealing tradeoffs between these mechanisms in microbial community resilience.

## Introduction

Structure-function relationships describe how the taxonomic composition and relative taxon abundances of a community determine its functional capacity. It is often assumed and commonly observed that changes in community structure drive changes in community function (Reed and Martiny, 2007; Graham et al., 2016; Galand et al., 2018). However, for microbial communities, structure-function relationships are not always apparent (Louca et al., 2018). For example, microbial functions can sometimes be better explained by differences in environmental conditions than in composition (e.g. (Langenheder et al., 2005; Louca et al., 2016b; Nelson et al., 2016; Ramírez-Flandes et al., 2019)), and dissimilar community compositions sometimes yield similar or identical functions (Bissett et al., 2011; Frossard et al., 2012; Purahong et al., 2014; Louca et al., 2016a; Jing et al., 2024). Thus, many microbial functions or activities seem to be uncoupled from the community structure of the resident microbes, at least some of the time.

Therefore, it remains a key pursuit to understand the contexts under which community structure determines microbial functions (Bier et al., 2015). Understanding structure-function relationships is especially critical in the face of intensifying climate change, which is altering the ranges and distributions of many organisms (Thomas, 2010; Hayden et al., 2012; Seabra et al., 2015; MacLean and Beissinger, 2017) and causing sudden, harsh disruption of communities and their functions after extreme events (Boucek and Rehage, 2014; Jurburg et al., 2017; Ummenhofer and Meehl, 2017; Maxwell et al., 2019; Sorensen et al., 2019; Nelson et al., 2022; Knight et al., 2024). Furthermore, microbial communities facilitate the biogeochemical cycles that feed back with climate change (Bardgett et al., 2008; Singh et al., 2010; Zhou et al., 2012; Wieder et al., 2013) and so predicting how structural changes will drive functional outcomes for these cycles is of urgency for response and adaptation (Wallenstein and Hall, 2012; Cavicchioli et al., 2019).

One potential explanation for the inconsistency in microbial community structure-function relationships may be due to high functional redundancy among different taxa within the community (Allison and Martiny, 2008). Functional redundancy (also known as “functional similarity”) describes the ability of coexisting taxa to perform similar functions (Walker, 1992; Lawton and Brown, 1994; Eisenhauer et al., 2023), therefore allowing for the possibility of a functional “replacement” of one taxon with another. Functional redundancy has often been considered in the ecological context of disturbance and ecosystem recovery as a mechanism to explain how the loss of one species may minimally affect overall ecosystem functioning due to the presence of a second species that can fill the same functional niche (Walker, 1992; Lawton and Brown, 1994; Rosenfeld, 2002). This forms the conceptual basis for the insurance, whereby a high diversity of populations within an ecosystem protects against functional changes or losses under environmental fluctuations or disturbance events (Yachi and Loreau, 1999). High functional redundancy is thus expected to promote ecosystem stability under changing conditions or disturbance as well as aid in ecosystem recovery (Naeem and Li, 1997; Awasthi et al., 2014; Tardy et al., 2014; Biggs et al., 2020), It is argued that few organisms are identical in their functional niches and can be replaced on a one-to-one basis due to their differences in response to environmental conditions, differences in their underlying cellular metabolism, and differences in process rates given the environment and the host cells (Schimel, 1995; Eisenhauer et al., 2023).

This applies especially to microorganisms, which have many metabolic processes shared across a vast range of taxonomic groups, particularly those related to carbon and energy processing and cellular biosynthesis, while other metabolisms, such as nitrogen or methane metabolism, are only found in select lineages (Schimel, 1995; Martiny et al., 2015). This variation in species-specificity can lead to differences in functional redundancy across metabolisms (Cheng et al., 2022). Thus, in the context of biogeochemical cycles important for climate change, there are some functional pathways, such as those related to heterotrophic respiration, that may be more likely to exhibit redundancy within a microbial community than others, like those related to denitrification. In reality, it may be more likely that there are species that overlap in some functions but not in others (Fischer and de Bello, 2023), and so no two species are completely functionally equivalent. Despite this limited evidence for 1:1 equivalency, community-level functions can remain stable when microbial functions are redundant, but not necessarily with identical environmental responses across taxa. In these cases, as environmental conditions change, organisms better adapted to the new conditions assume the functions of the less successful species (Fischer and de Bello, 2023). Without controlled studies, it remains difficult to determine an organism’s repertoire of environmental responses. However, from a genomic perspective, a microorganism’s ability to adapt to environmental change and modulate cellular functions is, in part, the outcome of its genomic investment in transcription factors (Cases et al., 2003; Santos et al., 2009). Transcription factors bind to DNA, either promoting or repressing transcription, thereby enabling an organism to control the activation and repression of operons and genes, and consequently, the activity of various pathways. Environmental variability can explain variation in transcription factor composition and abundance within bacterial genomes (Parter et al., 2007; Kostadinov et al., 2011; Angel et al., 2024). Organisms found in more environmentally dynamic habitats, such as soil, tend to encode more transcription factors in their genomes than those that are found in stable ones (Cases and de Lorenzo, 2005; Hoskisson and Rigali, 2009; Perez-Rueda et al., 2018).

Previously, we reported the strong response and recovery in the community structure of temperate soil bacterial microbiomes overlying the active coal seam fire in Centralia, Pennsylvania, USA (Barnett and Shade, 2024a). Since 1962, the underground Centralia fire has imposed an intense heat disturbance on the soils (Elick, 2011), leading to structurally distinct microbial communities across fire-affected and unaffected soils (Lee et al., 2017; Sorensen and Shade, 2020; Barnett and Shade, 2024a). As the fire advances along the coal seams, previously fire-affected soils recover in terms of community structure as temperatures return to ambient conditions (Barnett and Shade, 2024a). Here, we leverage the Centralia disturbance gradient to investigate the relationships between bacterial community structure, metagenome-informed functional redundancy, insurance potential, and resilience. Building on previous findings that showed a negative relationship between temperature and bacterial alpha diversity (Sorensen and Shade, 2020; Barnett and Shade, 2024a), we hypothesized that reference soils are more functionally redundant than fire-affected soils, with functional redundancy declining with increasing temperature and then increasing again after the disturbance subsided, thereby recovering diversity and supporting insurance.

## Methods

### Site description and soil sampling

Centralia, Pennsylvania, USA, lies above an anthracite seam that has been burning since 1962 (Elick, 2011). The fire heats the overlying soil to well above ambient temperature throughout the year (Elick, 2011; Barnett and Shade, 2024a) and imposes a strong homogenizing environmental filter at the temperature extremes (Sorensen and Lee, 2017). As the fire advances along the coal seam, the soils above the fire front warm, and the previously impacted soils cool. We have studied the Centralia fire gradient as a model to understand how environmental microbial communities respond to intense disturbance (Tobin-Janzen et al., 2005; Lee et al., 2017; Kearns and Shade, 2018; Sorensen et al., 2019; Sorensen and Shade, 2020; Barnett and Shade, 2024a, 2024b). In 2014, we established a series of sampling sites that represented different contemporary and historical exposures to fire, returning annually to collect new soil from these sites and quantify the microbial responses to the ongoing disturbance.

For the present study, we selected ten of these established sites, including three reference sites and seven fire-affected sites. These sites were sampled each October over seven years, from 2015 through 2021. Soil collection and processing for microbiome assessment have been described previously (Barnett and Shade, 2024a, 2024b). Additionally, air and soil temperatures, CO_2_ levels, gravimetric moisture levels, pH, and soil chemistry were measured (Barnett and Shade, 2024a).

### DNA and sequence processing

DNA extraction and processing was previously described (Barnett and Shade, 2024a, 2024b) and is detailed in supplemental. Similarly, we have previously detailed sequencing, initial sequence processing, and individual metagenome assembly (Barnett and Shade, 2024b) and in the supplemental. From the quality-controlled reads, we estimated metagenome coverages and community diversity using Nonpareil (Rodriguez-R and Konstantinidis, 2014) and average genome size using MicrobeCensus (Nayfach and Pollard, 2015). From the assemblies, only contigs 1 Kbp or longer were kept. Within these contigs, open reading frames (ORFs) were predicted using prodigal (Hyatt et al., 2010) version 2.6.3 as part of the PROKKA (Seemann, 2014) version 1.13 annotation pipeline.

ORFs were then mapped to KEGG orthologs (KO) using KofamScan version 1.3.0 with the KOfam database (Aramaki et al., 2020) downloaded April 24, 2023. ORFs were also mapped to carbohydrate active enzymes (CAZymes) from the dbCAN HMM database (Yin et al., 2012) version 11 using HMMER (http://hmmer.org) version 3.3.2. ORFs that were not annotated to a KO were dropped. Reads were mapped to the set of KEGG-annotated ORFs using BBMap (Bushnell, 2019).

Contigs were binned into metagenome assembled genomes (MAGs) using metaWRAP (Uritskiy et al., 2018) version 1.3.2. Binning was first performed for each sample individually with MetaBAT2 (Kang et al., 2019) version 2.12.1, MaxBin2 (Wu et al., 2016) version 2.2.6, and CONCOCT (Alneberg et al., 2014) version 1.0.0, and then refined into final sample-specific bins. Bin qualities were assessed with CheckM (Parks et al., 2015) version 1.0.12. These bins were then dereplicated across all samples into MAGs using dRep (Olm et al., 2017) version 3.4.0, keeping only MAGs with at least 50% completeness and less than 10% contamination. For dRep, the primary cluster ANI threshold was maintained at 0.9 while the secondary cluster ANI threshold was set at 0.99. MAG taxonomy was classified with GTDB-Tk (Chaumeil et al., 2022) version 2.1.1. Finally, reads from across all samples were mapped to the MAGs with BBMap. As a follow-up to the KEGG annotations to partially account for MAG incompleteness, we also predicted metabolic pathways in MAGs using gapseq (Zimmermann et al., 2021).

### Statistical Analyses

All statistical analyses were performed in R version 4.4.1, and code can be accessed via GitHub at https://github.com/ShadeLab/Centralia_metagenome_functional_redundancy_Barnett. We compared values between reference and fire-affected sites, regardless of year, using Wilcoxon rank sum tests. We evaluated trends across soil temperature with linear mixed effects (LME) models from the package *nlme* (Pinheiro et al., 2020) blocked by site ID to account for repeated sampling. When appropriate, p-values were adjusted for multiple comparisons using the Benjamini-Hochberg method (Benjamini and Hochberg, 1995). KO abundances were generated by first calculating the number of reads per thousand base pairs per million mapped reads (RPKM) for each ORF, then summing across all ORFs assigned to the same KO. MAG abundances were calculated as RPKM across the entire MAG length using just reads mapped to MAGs.

To examine broad variations in community functional potential across samples, we utilized beta-diversity metrics based on the abundance of KEGG orthologs within each sample. Between-sample diversity was calculated with the Bray-Curtis dissimilarity using RPKM for each KEGG orthologue. PCoA ordination was used to visualize this variation in metagenome content across samples. We compared variation in metagenome content to variation in community composition as measured by operational taxonomic unit relative abundances determined from amplicon sequencing of the same samples as previously described (Barnett and Shade, 2024a). The two measures were compared using both a Mantel test to correlate Bray-Curtis dissimilarity matrices and Procrustes analysis to correlate PCoAs. Mantel tests were performed with the function *mantel* and Procrustes analysis with functions *protest* and *procrustes,* all from the R package vegan (Oksanen et al., 2018). We performed a PERMANOVA with the R package vegan (Oksanen et al., 2018) to examine the extent to which fire classification and time explained variation in metagenome content. Constrained analysis of principal coordinates (CAP) was used to determine how measured soil properties further explained the variation in metagenome content as previously described. Edaphic factors considered were soil temperature (°C), concentrations (ppm) of calcium, iron, nitrate nitrogen, ammonium nitrogen, magnesium, phosphate, potassium, and arsenic, organic matter percentage, pH, and carbon dioxide release (ppm). Model selection was performed with the function ordistep from the *vegan* package. To examine inter-annual variation in community functional potential within each site using time-lag analysis. Time-lag analysis was compared between-timepoint Bray-Curtis dissimilarity and the square root of the timepoint difference.

Functional redundancy within samples was calculated using the methods reviewed in (Ricotta et al., 2016) based on MAGs. Functional redundancy was calculated as 1 –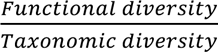, where functional diversity is Rao’s quadratic entropy and taxonomic diversity is the Simpson’s index. Rao’s quadratic entropy was calculated using the R package *adiv* (Pavoine, 2020) while Simpson’s index was calculated using the *vegan* package (Oksanen et al., 2018). Rao’s quadratic entropy considers functional differences across species as well as their abundances across samples. In applying Rao’s quadratic entropy to metagenomes, we calculated functional distance between MAGs as the Jaccard distance based on the presence and absence of KEGG orthologs detected in each MAG and defined MAG abundance as RPKM in each sample. Simpson’s index also utilized RPKM for MAG abundance.

Functional redundancy is expected to vary across types of functions (Cheng et al., 2022). We therefore focused on two major microbial processes that are important in soil: nitrogen cycling and carbohydrate metabolism. Nitrogen cycling pathways were defined by KEGG modules: nitrification (M00528), denitrification (M00529), dissimilatory/assimilatory nitrate reduction (M00530 and M00531), nitrogen fixation (M00175), anammox (M00973), and comammox (M00804). Carbohydrate metabolism was defined by all KEGG level C processes under level B “09101 Carbohydrate Metabolism”. We also examined glycoside hydrolases identified from the CAZy database (Drula et al., 2022; Zheng et al., 2023). We utilized both KEGG orthologs and glycoside hydrolase annotations for carbohydrate metabolism to cast a wider net beyond central metabolism, which comprises a large portion of pathways at the KEGG carbohydrate metabolism level. We also looked at the potential environmental breadth of the soil bacteria by examining the diversity of transcription factors across samples. For these functions, we utilized the full metagenome assemblies. We measured the number of genes recovered from nitrogen cycling pathways, carbohydrate degradation pathways, and transcription factor families relative to the number of genomes expected from the metagenomes (*i.e.,* genes per genome). The number of genomes expected from each metagenome sample was determined as the mean count of 35 ribosomal proteins and nine universal single-copy genes annotated within each sample (Parks et al., 2015). We assume recovery and assembly of universal single-copy genes is correlated with other genes within a sample. By adjusting the gene counts for the number of universal single-copy genes (*i.e.,* genes per genome), we account for differences in gene recovery due to coverage and assembly differences between samples. This method is derived from methods to normalize gene abundances (Shaomei et al., 2015; Nayfach and Pollard, 2016). In evidence, the number of universal single-copy genes recovered was strongly correlated to metagenome coverage as measured by nonpareil (linear mixed effects regression: R^2^ = 0.797, p-value < 0.001).

## Results

### Soil microbiome functional genes were relatively stable over the disturbance gradient

First, we asked whether the metagenomes exhibited similar responses to the fire gradient as we observed for the amplicon profiles. Read-based microbial diversity (*i.e.,* Nonpareil) followed the same general patterns previously observed for 16S rRNA gene amplicon profiling (Barnett and Shade, 2024a): functional gene diversity was higher in reference than in fire-affected soils and recovered over time as the fire-affected soils cooled (Fig. S1A). Also, read-based estimated average genome size was larger in reference soils as compared to fire-affected soils, in agreement with previous results from a single-year metagenome analysis that reported ecological selection for smaller cells and genome sizes with higher temperatures (Sorensen et al., 2019) Fig. S1B). The average estimated genome size also increased in the fire-affected soils over time, as cooling occurred.

Metagenome and amplicon profiles were concordant (Procrustes analysis: correlation = 0.802, m_12_^2^ = 0.357, p-value = 0.001; Fig. S2 and Fig. S3). From PERMANOVA and CAP, the same environmental variables (e.g., time, temperature, pH, etc.) that explained variance in the amplicon profiles were also explanatory for the metagenomes (see Supplemental Information; Table S1; Figs. 1A, S3, and S4). Thus, functional gene and amplicon profiles were generally aligned in their responses to disturbance in both alpha and beta diversity.

**Figure 1:**
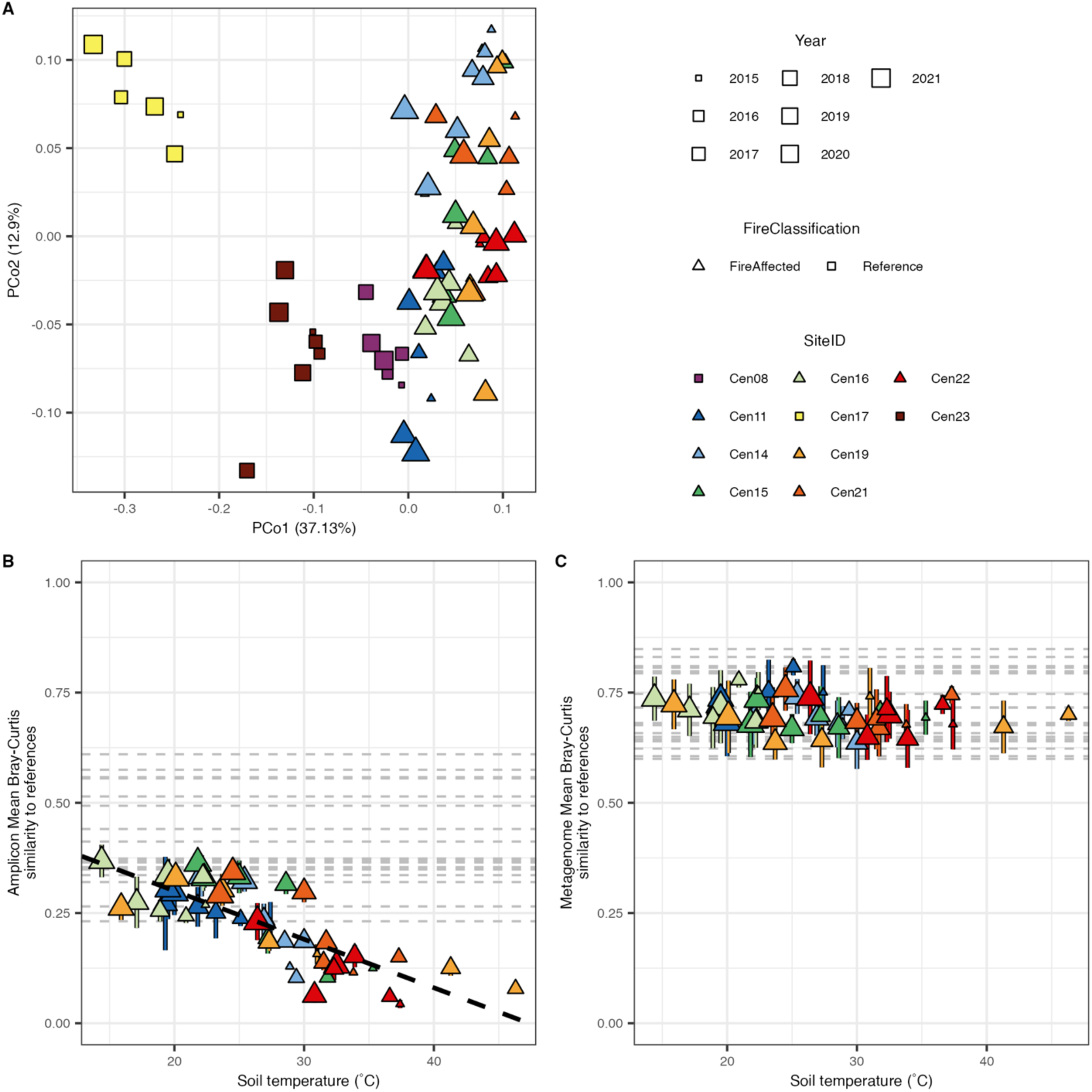
Variation in functional gene (metagenome) and taxonomic (16S rRNA amplicon) profiles across samples. A) PCoA of metagenome profiles based on Bray-Curtis dissimilarity KEGG orthologues across samples. B) Amplicon-based community composition shows relatively higher resilience, whereby OTU Bray-Curtis similarity between fire-affected sites and matching reference sites increases as disturbed soils cool. C) There is consistent and relatively high functional gene (KEGG orthologues) Bray-Curtis similarity between fire-affected sites and matching reference sites, even as disturbed soils cool off.

**Figure 2:**
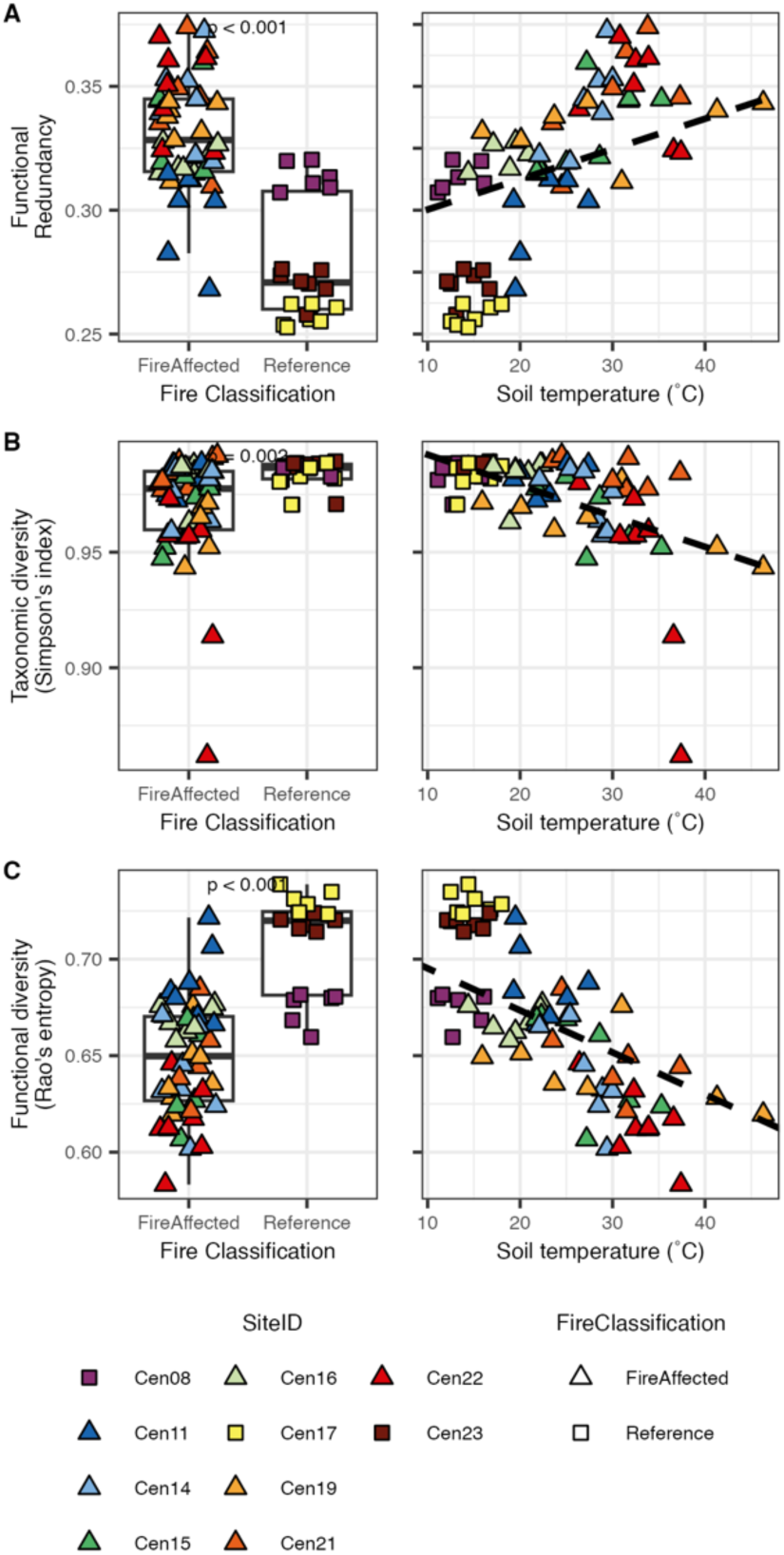
Functional redundancy as measured by the relationship between taxonomic diversity (Simpson’s index) and functional diversity (Rao’s entropy). A) Functional redundancy is higher in fire-affected soils than in reference soils and decreases as soil temperature decreases. B) Taxonomic diversity is higher in reference than in fire-affected soils and increases as soils cool off. C) Functional diversity is higher in reference than in fire-affected soils and increases as soils cool off.

**Figure 3:**
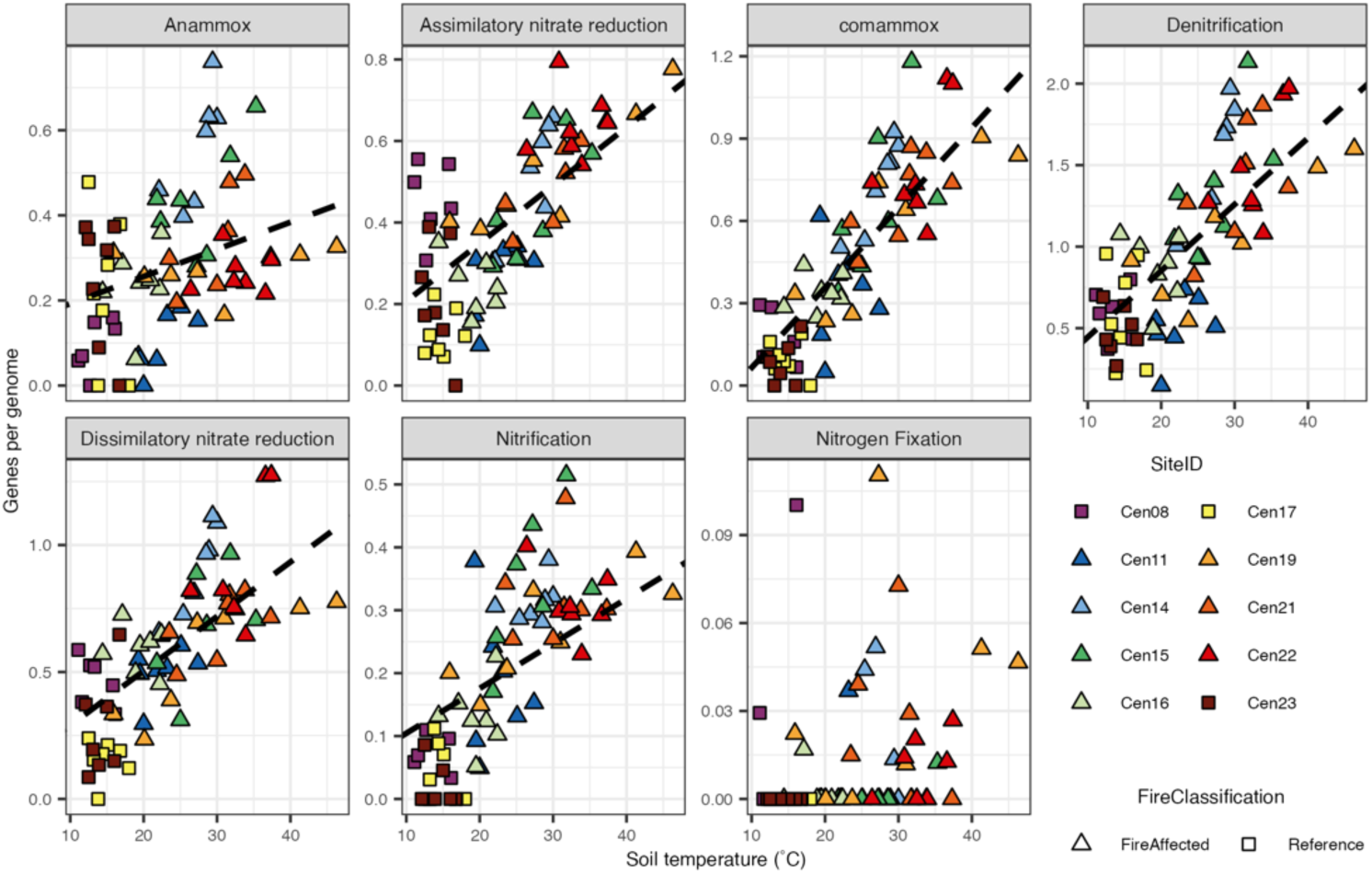
Per-genome investment in Nitrogen cycling pathways. Pathway investment is calculated as the total number of KEGG orthologs within each pathway detected across open reading frames within each sample, divided by the median number of single-copy genes also identified.

Resilience can be expressed as a rate of change from a disturbed towards a recovered condition (Barnett and Shade, 2024a). Thus, we calculated resilience as the change in community similarity for taxonomic structure (amplicon profiles) and functional gene structure (metagenome profiles) over the disturbance gradient. There was an inverse relationship between temperature (as a proxy for disturbance intensity) and recovery in community structure (slope = -0.011, p-value < 0.001; Figure 1B), confirming a resilient composition with cooling. However, the functional gene structure remained relatively stable over the disturbance gradient (p-value > 0.05; Figure 1C).

### Selective disturbance strengthened functional redundancy

From the assembled contigs, we produced 1212 unique medium- and high-quality MAGs, representing 23 archaeal MAGs across two phyla and 1189 bacterial MAGs across 26 phyla. We found that this dataset generally represented the major lineages and reproduced the dynamics in beta-diversity observed for 16S rRNA amplicon profiling (see Supplemental results, Figs. S6-S9). Thus, we calculated functional redundancy based on the differences in MAGs’ gene content and abundances across soils.

Functional redundancy was higher in the disturbed soils compared to the reference soils (Wilcoxon test: W = 61, p-value < 0.001, Fig. 2). Over time, and as the soils cooled, functional redundancy in the fire-affected soils decreased, though it did not ultimately achieve the same values as the reference soils (LME: slope = 0.001, p-value = 0.001; Fig. 2A). When considering the functional redundancy metric split into its components, Simpson’s diversity index (*i.e.,* taxonomic diversity) and Rao’s quadratic entropy (*i.e.,* functional diversity), there were further distinctions between our disturbed and undisturbed soils. Both taxonomic and functional diversity are generally higher in the reference soils than the fire-affected soils (Wilcoxon tests; Simpson’s index: W = 725, p-value = 0.002; Rao’s quadratic entropy: W = 916, p-value < 0.001) and increased as soils cooled (LME; Simpson’s index: slope = -0.001, p-value < 0.001; Rao’s quadratic entropy: slope = -0.002, p-value < 0.001; Fig. 2B and C). Similar results are obtained when using the presence/absence of metabolic pathways predicted from the MAGs with gapseq (Supplemental results; Fig. S10).

### Bacteria invest differently in specific functions with disturbance and recovery

We examined the disturbance responses of several microbial functions that are important in soil nitrogen cycling and carbohydrate processing. For these functions, we utilized the full metagenome assemblies. We determined the number of genes recovered relative to the number of genomes expected from the metagenomes (*i.e.,* genes-per-genome, see methods). As expected, there was slightly higher metagenome coverage in fire-affected soils due to their relatively lower community alpha diversity. The mean number of ribosomal proteins and universal single-copy genes was higher in fire-affected than reference soils and decreased with decreasing temperature (LME: slope = 1.594, p-value < 0.001; Fig. S11).

We first considered genes related to processes within the nitrogen cycle (*i.e.,* nitrification, denitrification, dissimilatory/assimilatory nitrate reduction, nitrogen fixation, anammox, and comammox), which generally are reported to be more specialized functions that range in redundancy among bacterial taxa (Nelson et al., 2016). For all processes except nitrogen fixation, the number of genes per genome decreased with decreasing temperature (Fig. 3, Table S2). Nitrogen fixation genes were rare and had no relationship with temperature.

We next looked at the genes related to carbohydrate metabolism, examining both genes within various carbohydrate processes (*i.e.,* KEGG level B “09101 Carbohydrate Metabolism”) and glycoside hydrolases from the CAZyme database. Again, for most processes under the KEGG level C level, we found that the number of genes per genome decreased with decreasing temperature (Fig. 4, Table S2). Both CAZyme and glycoside hydrolase genes per genome were significantly higher in fire-affected soils than in the reference soil (Fig. S12). However, neither had a statistically supported linear relationship with temperature.

**Figure 4:**
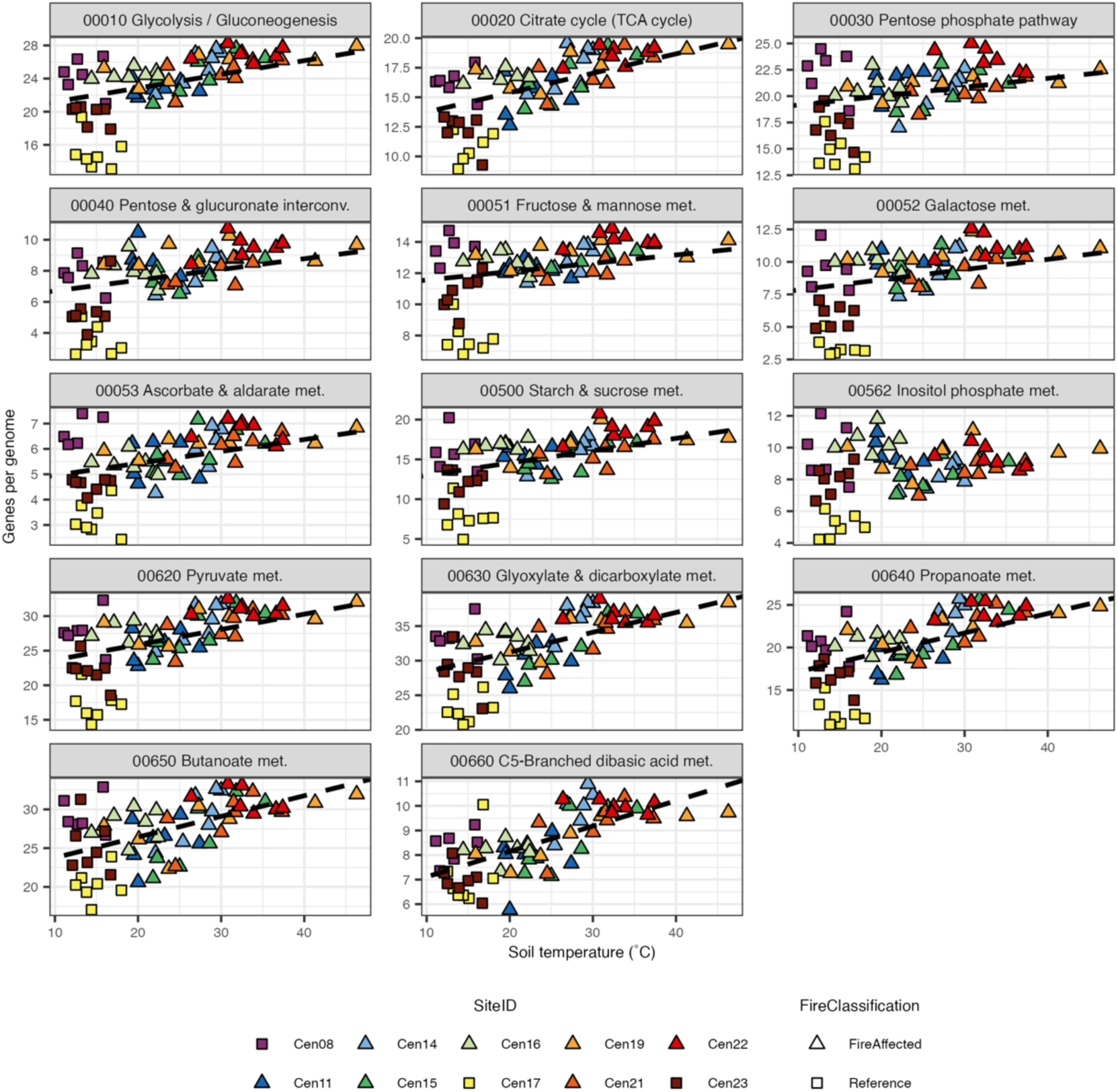
Per-genome investment in carbohydrate metabolism pathways. Pathway investment is calculated as the total number of KEGG orthologs within each pathway detected across open reading frames within each sample, divided by the median number of single-copy genes also identified.

### The relationship between transcription factor investment and temperature varies by gene family

We finally aimed to examine the potential environmental breadth of the microorganisms in soils along the disturbance gradient. As transcription factor diversity has been shown to relate to environmental breadth, we quantified the transcription factor genes per genome and related these values to soil temperature. As before, we first examined the number of transcription factor genes recovered relative to the mean ribosomal protein and single-copy genes (*i.e.,* genes per genome). This analysis was done separately for each family of transcription factor as defined in KEGG. Of the 44 transcription factor families with genes identified in the metagenome dataset, 20 showed significant relationships between genes per genome and soil temperature (Fig. S13; Table S2). Notably, the patterns differed by transcription factor family. Specifically, while ArsR, BirA, CsoR, DtxR, GntR, HTH-type, IclR, LacI, LuxR, MerR, ParB, PucR, Rrf2, and Sigma-54 dependent families demonstrated fewer genes per genome as soil temperature decreased, genes per genome for AraC, BlaI, CarD, Lrp/AsnC, and PadR families increased as soil temperature decreased.

We next examined the evenness of transcription factor orthologs based on RPKM abundance. Overall transcription factor evenness was higher in reference soils than in fire-affected soils (Wilcoxon test: W = 114, p-value < 0.001). However, no significant relationship with soil temperature was found (LME: p-value = 0.178). We then looked specifically at different families of transcription factors as before, excluding families with fewer than five distinct orthologues recovered across all samples. Five transcription factor families had statistically significant relationships between orthologue evenness and temperature, including AraC, Fur, GntR, HTH-type, and Rrf2 families (Figure 5, Table S3). In all cases, evenness increased as soil temperature decreased.

**Figure 5:**
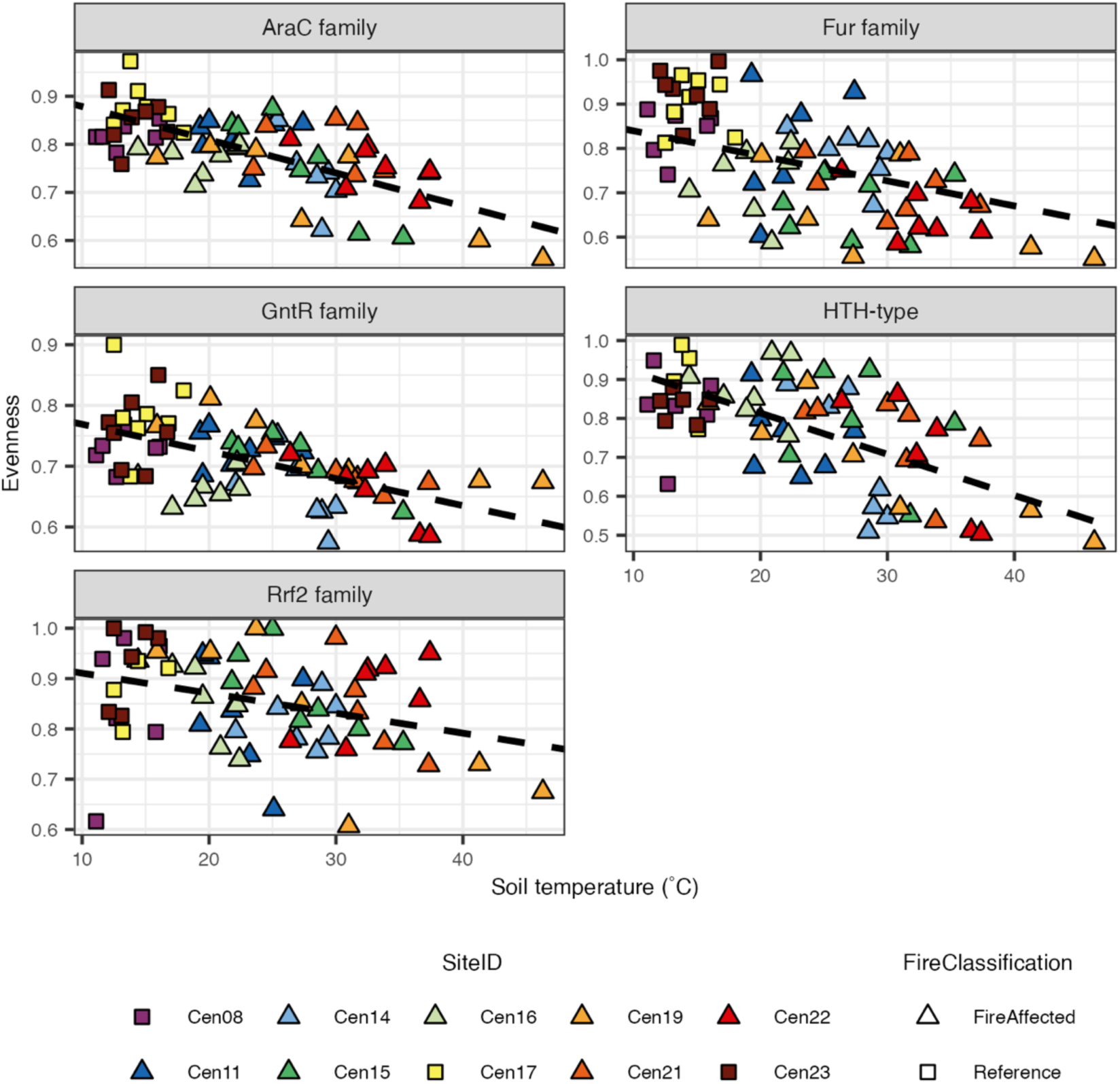
Evenness of five transcription factor families that vary across soil temperature. Evenness is calculated as Pielou’s Evenness across KEGG orthologs within each family. Note that the evenness axis varies across transcription factor families. The HTH-type family includes helix-turn-helix domain-containing genes not already included in another family.

## Discussion

The underground coal mine fire in Centralia, PA, offers an interesting case study to examine the relationship between bacterial community structure, functional potential, and community resilience in the context of a long-term and ongoing anthropogenic disturbance. Incorporating all identified KEGG orthologs into a measure of functional gene similarity, we observed relatively low dynamics in functional gene composition over time with high similarity between disturbed and undisturbed soils, unlike the strong dynamics observed for community structure using either amplicons (Barnett and Shade, 2024a) or MAGs. This seeming incongruence in structure with functional potential suggests redundancy is evident across these soils.

Our original hypothesis was that undisturbed communities are more diverse and thus also more functionally redundant and supportive of insurance than disturbed communities, but this was not precisely supported. The higher functional redundancy observed in the heated soils relative to reference soils was the outcome of a homogenizing selection imposed by the long-term heating disturbance (Lee et al., 2017; Barnett and Shade, 2024a), which reduced both taxonomic and functional diversity. Thus, the surviving taxa are relatively more genomically similar and thus more functionally redundant with each other. This observed dynamic is a logical outcome of functional redundancy, as illustrated by Fischer and de Bello (Fischer and de Bello, 2023), where redundancy in the original community is expanded when there is a loss of functions and metabolisms with the loss of community members, such as rare or unfit taxa. This process of gradual selection, which results in increased functional redundancy, is most evident in relatively widespread microbial metabolisms, such as carbon and nitrogen processing, that tend to be maintained under disturbance (Souza et al., 2015; Nelson et al., 2016). There was greater per-genome investment in pathways associated with both metabolisms in fire-affected soils relative to reference soils. We further confirmed the decrease in average genome size with increasing temperature (Sorensen et al., 2019), indicating a selection for smaller (potentially more efficient or stable) genomes (Sabath et al., 2013) and thus possibly the concurring loss of rare functions that are not favorable in the context of the strong selection (Ruhl et al., 2022). Notably, increased per-genome investment in carbon and nitrogen metabolisms does not necessarily indicate increased activity of these pathways because functional gene presence suggests only functional potential and cannot reveal in situ functions directly (Helbling et al., 2012). However, the relationships between these functional genes and disturbance are indicative of environmental selection.

Contrastingly, while we found evidence pointing to selection within environmental response mechanisms, this selection does not have a consistent relationship to temperature. Specifically, we observed variable per-genome investment patterns across diverse families of transcription factors, with some increasing and others decreasing in response to soil temperature. As transcription factors are important indicators of microbial responses to environmental change (Cases et al., 2003; Santos et al., 2009), the observed shift in transcription factor investment suggests a decreased ability of fire-affected community members to respond to subsequent different or compounded environmental disturbances. This proposed selection in transcription factors may explain patterns of environmental responsiveness in soil microbial communities under multiple and successive stressors (Philippot et al., 2021). For example, soils that were heated resulted in a successional microbial community that, when heated a second time, underwent a similar community shift as observed when heating previously undisturbed soils (Jurburg et al., 2017). However, when those initially heated soils underwent a freezing treatment, the community shifted in a unique way (Jurburg et al., 2017). The conclusion was that there was selection for a heating response following a heat disturbance, thereby maintaining the community’s response to heat in the successional community. However, there was no selection for a freezing response; thus, the original responsiveness to freezing was altered.

As described by Fischer and de Bello (Fischer and de Bello, 2023), functional redundancy in the context of the insurance hypothesis may require disturbance-resistant taxa to take over functions from disturbance-sensitive taxa. Thus, if the initial disturbance of soil heating reduces the resistance of the surviving community to a different or compounding disturbance (Philippot et al., 2021), the subsequent disturbance may not be tolerated by the initially surviving taxa, with a simultaneous decrease in overall insurance. Thus, it is not just functional redundancy that provides community insurance against disturbance, but also the repertoire of environmental responsiveness retained by the community’s members. One proxy for environmental responsiveness among bacterial communities may be the evenness of transcription factors. It has been proposed that community evenness, in conjunction with functional redundancy, can serve as an indicator of community functional stability in response to stress (Wittebolle et al., 2009). Low evenness may lead to changes in community function under disturbance, even if functional redundancy is generally high, due to loss of the dominant taxa and severe shifts in community structure (Fonseca and Ganade, 2001). Generally, there was a negative relationship between microbiome transcription factor evenness and soil temperature, indicating that, in hotter soils, certain transcription factors, and thus potential environmental responsiveness, are more heavily enriched over others. Given this observation, undisturbed soils may be better insured against further diverse disturbances than disturbed soils due to their relatively even representation of transcription factors.

The transcription factors showing negative trends are highly diverse in terms of their known functional regulations (Rodionov, 2007). Both AraC and GntR family genes are involved in a wide array of functions, including carbon and nitrogen metabolism, stress response, and even virulence factors (Ibarra et al., 2008; Hoskisson and Rigali, 2009). The strong link between the AraC and GntR families and the control of catabolism of various carbon and nitrogen sources is particularly intriguing in this environment, where fire-related disturbances may be altering the organic matter available to the microbes (Tobin-Janzen et al., 2005; Janzen and Tobin-Janzen, 2008). Fur family transcription factors are commonly associated with metalloregulation, especially iron, but including zinc, manganese, and nickel (Lee and Helmann, 2007). While we did not measure or observe significant changes in concentrations of all these metals across Centralia soils, metal availability can be significantly affected by coal combustion (Mandal and Sengupta, 2006) and pH, which we have shown decreases in Centralia soils after heating, likely due to acidification from coal combustion (Barnett and Shade, 2024a). Rrf2 family transcription factors are known for their use in response to stress, particularly iron and nitric oxide (Banerjee et al., 2024), again both potentially associated with coal combustion (Mitchell and Tarbell, 1982; Mandal and Sengupta, 2006). Thus, the decreased evenness of these transcription factor families under soil heating indicates a winnowing of the environmental responsiveness of these communities due to the strong selection by the disturbance.

As is true of all MAG-based analyses, this study is reliant on the completeness and diversity of the recovered MAGs. Since the functional redundancy metric used here utilizes differences in MAG gene composition to measure functional variation across genomes, incomplete genome recovery may lead to artificially greater functional dissimilarity across genomes. However, we used only medium- and high-quality MAGs, and reduced any bias due to genome incompleteness by dereplicating MAGs across all samples. Furthermore, we also tested functional redundancy at the pathway level, rather than individual KEGG orthologue level, using gapseq. Gapseq predicts pathway presence in genomes even with missing components (Zimmermann et al., 2021). Results from gapseq pathways were similar to those of the KEGG orthologs alone. Furthermore, we provide evidence that the MAGs assembled were representative of the taxonomic diversity we observed previously using 16S rRNA gene amplicon sequencing, and thus useful for these functional redundancy measures. Combining both MAG-based and unbinned assembly-based analyses further provided a more complete picture of the variation in functional potentials across the sites. Future development on community-aggregated trait-based approaches may yield further refined methods for measuring functional diversity and redundancy from metagenomic sequencing (Finn, 2024).

Only recently have methodological and technological innovations allowed us to investigate functional redundancy in highly diverse microbial communities (Ramond et al., 2025). Our study exemplifies the effort to understand better microbial community functional redundancy in the context of anthropogenic disturbance, utilizing metagenomic sequencing and both assembly- and MAG-based analyses. We demonstrated here that methods developed for use in macroecology contexts (Ricotta et al., 2016) apply to microorganisms and that controlling for species richness is important, particularly where species loss due to environmental disturbance is expected. Our findings add to the growing body of evidence that functional redundancy plays a major role in microbial community response to large-scale environmental change (Allison and Martiny, 2008; Ramond et al., 2025). More specifically, our findings show a tradeoff in functional redundancy and insurance potential for resilience across complex or compounded disturbances.

Disturbance-impacted ecosystems like Centralia can offer valuable real-world opportunities to uncover dynamics in functional redundancy and related measures (Ladau and Eloe-Fadrosh, 2019). Such experimentation and development of standardized approaches to measure functional diversity and functional redundancy, including the ones used here, contribute to our understanding of microbiome dynamics in the face of climate change and other environmental perturbations (Shade, 2023; Ramond et al., 2025).

## Data Availability Statement

Raw reads, assembled contigs at least 1,000 bp long, and MAG sequences are available through NCBI in the SRA and genome databases, under BioProject PRJNA973689. Code for read processing and assembly can be accessed via GitHub at https://github.com/ShadeLab/Centralia_7year_metagenome_processing_Barnett_2024, while annotation, MAG processing, and analysis code can be accessed via GitHub at https://github.com/ShadeLab/Centralia_metagenome_functional_redundancy_Barnett.

## Acknowledgments

This work was supported by the U.S. National Science Foundation CAREER award #1749544 to AS.

## Supplemental file

Word document including supplemental Methods, Results, and Supplemental Figures S1-S13, and Supplemental Tables S1-S3.

