## Supplemental Material for "Disturbance increases soil microbiome functional redundancy but decreases capacity for insurance via winnowed environmental responsiveness"

### Supplemental Materials and Methods

*DNA and sequence processing*

DNA was extracted from soils using a standardized phenol-chloroform protocol (Griffiths et al., 2000) which was modified to use 0.1 mm zircondium bead BeadBug homogenizer tubes (#Z763764; Benchmark Scientific, Sayreville, NJ, USA) in the bead beating step. Illumina untargeted metagenomic DNA libraries were prepared and sequenced by the Research Technology Support Facility at Michigan State University (East Lansing, MI, USA) following standard protocols. Libraries were barcoded using dual indexes prior to pooling with the ThruPLEX DNA-Seq kit (Takara Bio Inc., Kusatsu, Japan) and sequenced in two lanes of a NovaSeq S4 flow cell in 2 x 150 bp paired-end format with the NovaSeq 6000 (Illumina, San Diego, CA, USA).

Demultiplexed metagenome sequencing sets were processed using a custom workflow based on the Joint Genome Institute SOP as previously described (Barnett and Shade, 2024b). From quality controlled reads, we estimated microbiome diversity using Nonpareil (Rodriguez-R and Konstantinidis, 2014) and average genome size using

MicrobeCensus (Nayfach and Pollard, 2015). Quality controlled and error corrected reads were assembled into contigs separately for each sample with SPAdes (Nurk et al., 2017) version 3.15.5 in metaSPAdes mode without internal read error correction. Contigs under 1,000 bp were discarded. Details of metagenome sequencing have been previously reported as a data resource (Barnett and Shade, 2024b). In summary, combined libraries from technical replicates produced between 47-172 million quality-filtered reads per sample. After metagenome assembly per sample, we recovered between 18,000-192,000 contigs per sample longer than 1 Kbp. These contigs had N50 values ranging from 1,458-11,958 bp.

### Supplemental Results

*Soil microbiome functional potential was less dynamic than taxonomic structure*

We examined the whole assembled metagenomes over time to observe broad shifts in community functional potential as the fire affected sites cooled off. PCoA Ordinations clearly demonstrate clustering of community gene profiles by site and fire classification (Fig 1A). PERMANOVA (Table S1) further indicates that fire classification and, to a lesser extent, time both explain significant portions of the variation in community gene profiles (fire classification: $R^2$ = 0.261, p-value =0.003; time: $R^2$ = 0.062, p-value = 0.002). As expected, constrained analysis of principal coordinates (CAP), demonstrated that multiple soil properties explained variation in the community gene profiles (45.652% overall). Most notably, temperature, and associated factors ($CO_2$ and soil organic matter) clearly distinguished the reference from fire affected soils while pH and associated factors (Ca) distinguished sites within both fire classes (Fig. S4).

When we look at inter-annual variation in community gene profiles within each site using time-lag analysis, There was relatively consistent variation across all year spans in reference sites but increasing variation across timepoints as year difference increases in fire affected sites (Fig. S5A). Notably, inter-annual variation is lower for smaller year ranges for fire affected sites than for reference sites, though the overall range of dissimilarity is similar between the two fire classifications. The slopes of the time-lag comparisons across fire affected sites increases as maximum soil temperature (*i.e.,* disturbance intensity) increases (Fig. S5B).

*MAGs broadly represented diversity observed in amplicon based analysis*

When mapping metagenome reads to MAGs, the mapped reads accounted for between 11.25-88.33% of reads per sample. Fewer reads tended to map to MAGs in reference soils than fire affected soils and decreased as soil temperature decreased (Fig. S6). Taxonomic assignment using GTDB-Tk identified. Generally, bacterial phylum-level composition from MAGs was comparable to what was observed with the amplicon sequencing (Fig. S7). For example, there were relatively more *Chloroflexi* in the fire-affected soils than the reference soils while Actinobacteria are more relatively abundant in the cooler reference soils.

Structure of bacterial communities determined using MAGs was further found to be strongly correlated those using OTUs from amplicon sequencing with a Mantel test, based on Bray-Curtis dissimilarity (Mantel: $r = 0.880$, p-value < 0.001), and a Procrustes analysis, based on Bray-Curtis dissimilarity and Principal coordinate analysis (Procrustes: correlation = 0.956, m12 = 0.087, p-value = 0.001; Fig. S4; Figs. S8 and S9).

*Similar redundancy patters observed with pathways rather than KEGG orthologues*

As MAGs are almost always incomplete, with our minimum cutoff of 50% completeness, we are likely missing KEGG orthologues in many our MAGs. To control for this in some way, we also ran our functional redundancy analysis using pathways identified by gapseq. Since gapseq includes incomplete pathways, random loss of genes due to limited completeness should be less of a factor. We found similar results as using KEGG orthologues. Functional redundancy was higher in the fire-affected soils compared to the reference soils (Wilcoxon test: $W = 197$, p-value < 0.001). Over time, and as the soils cooled, functional redundancy in the fire affected soils decreased (LME: slope = 0.001, p-value = 0.005; Fig. S10A). Both taxonomic and functional diversity are generally higher in the reference soils than the fire affected soils (Wilcoxon tests; Simpson's index: $W = 725$, p-value = 0.002; Rao's quadratic entropy: $W = 853$, p-value < 0.001) and increases as soils cool off (LME; Simpson's index: slope = -0.001, p-value < 0.001; Rao's quadratic entropy: slope = -0.002, p-value < 0.001 ; Fig. 2B and C).

### Supplemental Tables

| Data set | Factor | Df | SumOfSqs | R2 | F | F Pr(>F) |
|---|---|---|---|---|---|---|
| **Amplicon** | Fire classification | 1 | 3.0389 | 0.16211 | 12.3303 | 0.001 |
| | Year | 6 | 1.3972 | 0.07453 | 0.9448 | 0.001 |
| | Fire classification: Year | 6 | 0.7546 | 0.04026 | 0.5103 | 0.001 |
| | Residual | 55 | 13.5554 | 0.72310 | NA | NA |
| | Total | 68 | 18.7461 | 1.00000 | NA | NA |
| **Metagenome** | Fire classification | 1 | 0.58059 | 0.26090 | 23.2830 | 0.003 |
| | Year | 6 | 0.13903 | 0.06248 | 0.9293 | 0.002 |
| | Fire classification: Year | 6 | 0.13425 | 0.06033 | 0.8973 | 0.112 |
| | Residual | 55 | 1.37150 | 0.61630 | NA | NA |
| | Total | 68 | 2.22538 | 1.00000 | NA | NA |

**Table S1:** PERMANOVA output for OTUs and KEGG orthologs.

| Metabolism | Pathway/Family | Slope | p-value |
|---|---|---|---|
| **Carbohydrate metabolism** | Glycolysis / Gluconeogenesis | 0.166 | < 0.001 |
| | Citrate cycle (TCA cycle) | 0.164 | < 0.001 |
| | Pentose phosphate pathway | 0.084 | 0.033 |
| | Pentose & glucuronate interconversions | 0.069 | 0.010 |
| | Fructose & mannose metabolism | 0.054 | 0.015 |
| | Galactose metabolism | 0.078 | 0.007 |
| | Ascorbate & aldarate metabolism | 0.047 | 0.007 |
| | Starch & sucrose metabolism | 0.155 | < 0.001 |
| | Pyruvate metabolism | 0.221 | < 0.001 |
| | Glyoxylate & dicarboxylate metabolism | 0.287 | < 0.001 |
| | Propanoate metabolism | 0.228 | < 0.001 |
| | Butanoate metabolism | 0.266 | < 0.001 |
| | C5-Branched dibasic acid metabolism | 0.103 | < 0.001 |
| **Nitrogen metabolism** | Assimilatory nitrate reduction | 0.014 | < 0.001 |
| | Dissimilatory nitrate reduction | 0.021 | < 0.001 |
| | Denitrification | 0.041 | < 0.001 |
| | Nitrification | 0.007 | < 0.001 |
| | Comammox | 0.029 | < 0.001 |
| | Anammox | 0.006 | 0.023 |
| **Transcription factors** | AraC family | -0.052 | < 0.001 |
| | ArsR family | 0.021 | 0.005 |
| | GntR family | 0.056 | 0.021 |
| | IclR family | 0.017 | 0.005 |
| | LacI family | 0.061 | 0.002 |
| | Lrp/AsnC family | -0.030 | 0.002 |
| | LuxR family | 0.031 | 0.019 |
| | MerR family | 0.026 | 0.002 |
| | PadR family | -0.170 | < 0.001 |
| | Rrf2 family | 0.009 | < 0.001 |
| | DtxR family | 0.016 | < 0.001 |
| | BlaI family | -0.052 | < 0.001 |
| | BirA family | 0.008 | 0.001 |
| | HTH-type | 0.013 | 0.001 |
| | Sigma 54 dependent | 0.002 | 0.016 |
| | CsoR family | 0.008 | 0.016 |
| | PucR family | 0.010 | 0.006 |
| | BolA family | -0.005 | 0.001 |
| | ParB family | 0.025 | < 0.001 |
| | CarD family | -0.011 | 0.001 |

**Table S2:** Linear mixed effects model coefficients for per-genome investments in carbohydrate and nitrogen metabolism pathways and transcription factor families. P-value is adjusted for multiple comparisons using the Benjamini & Hochberg method (carbohydrate metabolism n = 14, nitrogen metabolism n = 7, and transcription factors n = 20). Only pathways and transcription factor families with p-values below the significance cutoff of 0.05 were included.

| Family | Slope | p-value |
| --- | --- | --- |
| AraC | -0.007 | < 0.001 |
| Fur | -0.006 | 0.008 |
| GntR | -0.004 | < 0.001 |
| HTH-type | -0.011 | < 0.001 |
| Rrf2 | -0.004 | 0.016 |

**Table S3:** Linear mixed effects model coefficients for evenness within transcription factor families. P-value is adjusted for multiple comparisons using the Benjamini & Hochberg method (n = 17). Only pathways and transcription factor families with p-values below the significance cutoff of 0.05 were included.

### Supplemental figures

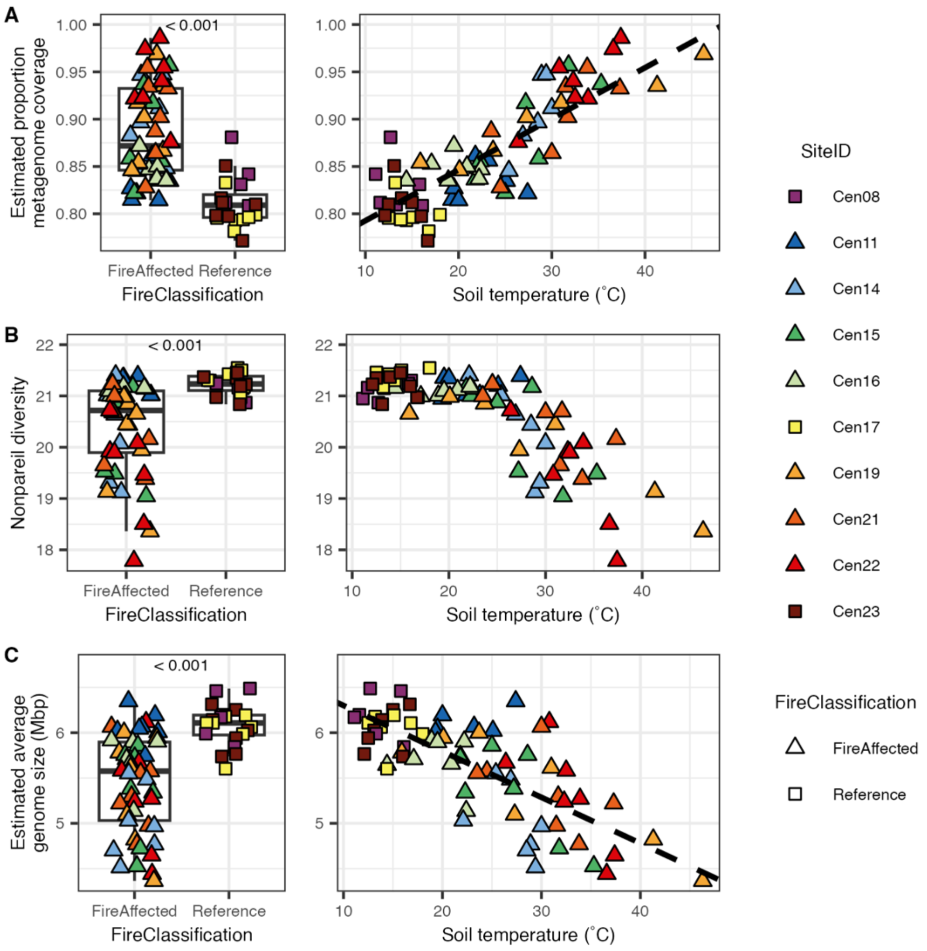


**Figure S1:** Read based community analyses. For all we examined values across fire classification and across soil temperature. A) Estimated proportion of metagenome covered by reads based on nonpareil. B) Estimated community diversity as measured by nonpareil. C) Estimated average gnome size as measured by microbecensus. For comparisons across fire classification p-value is indicated above the points (Wilcoxon test). For relationship to temperature, dashed lines indicate significant linear relationship (linear mixed effects model)

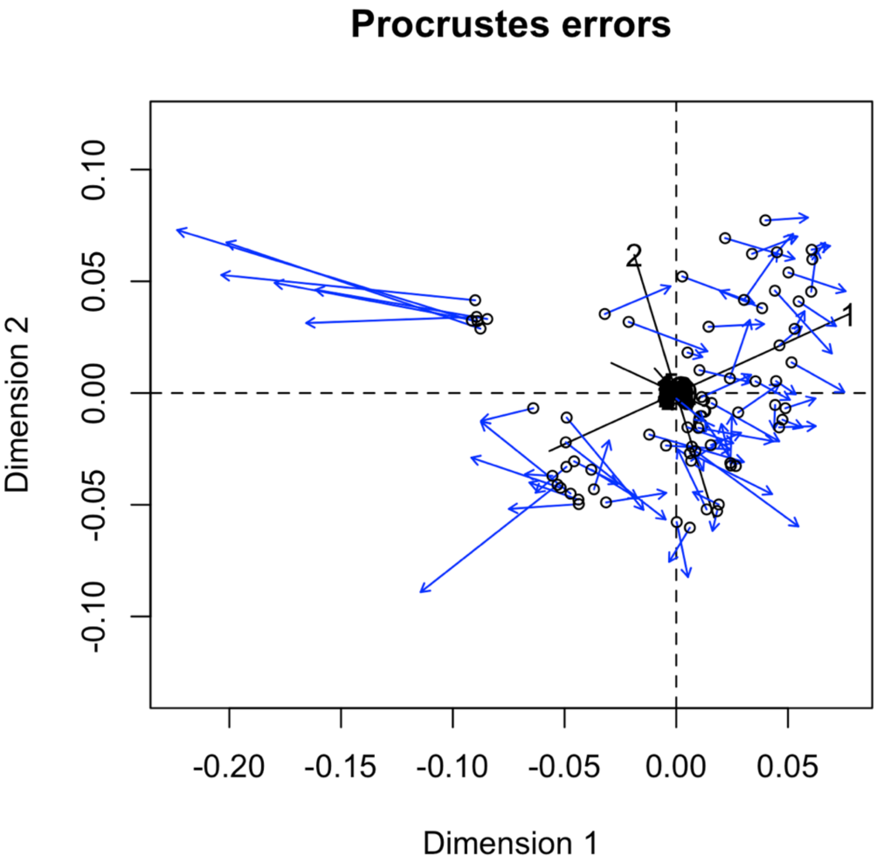


**Figure S2:** Plot of Procrustes errors between metagenome (KEGG orthologue) and amplicon (OTU) community structures. Ordinations used were based on Bray-Curtis dissimilarity and PCoA.

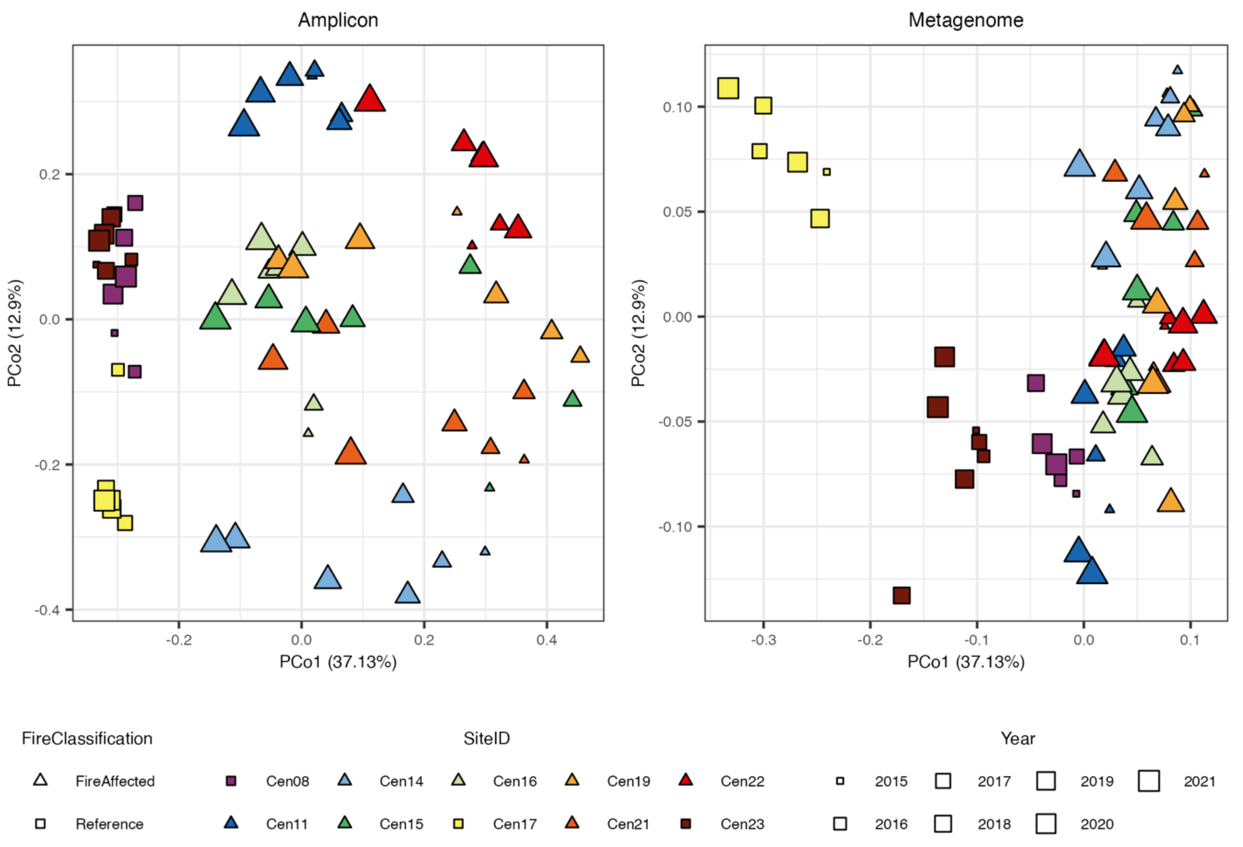


**Figure S3:** PCoA of Amplicon (OTU) and metagenome (KEGG orthologue) based community structures. Both utilized Bray-Curtis dissimilarity.

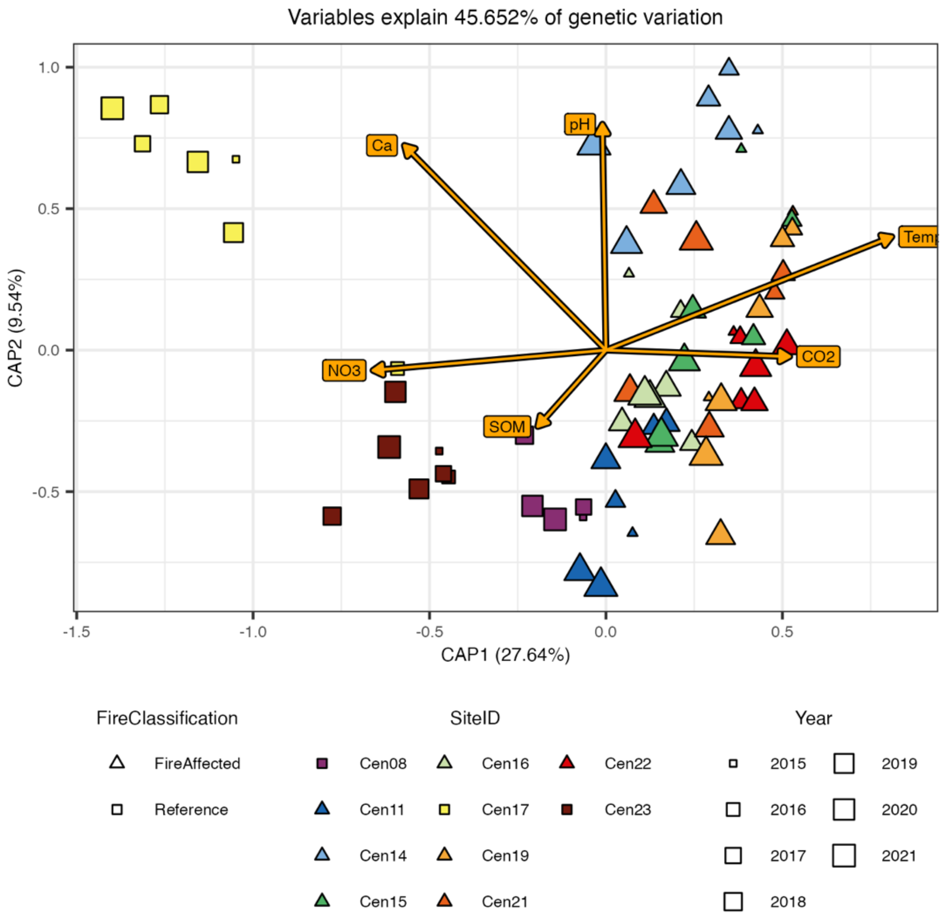


**Figure S4:** CAP analysis indicating metagenome structure (KEGG orthologue) variation explained by measured environmental and edaphic factors.

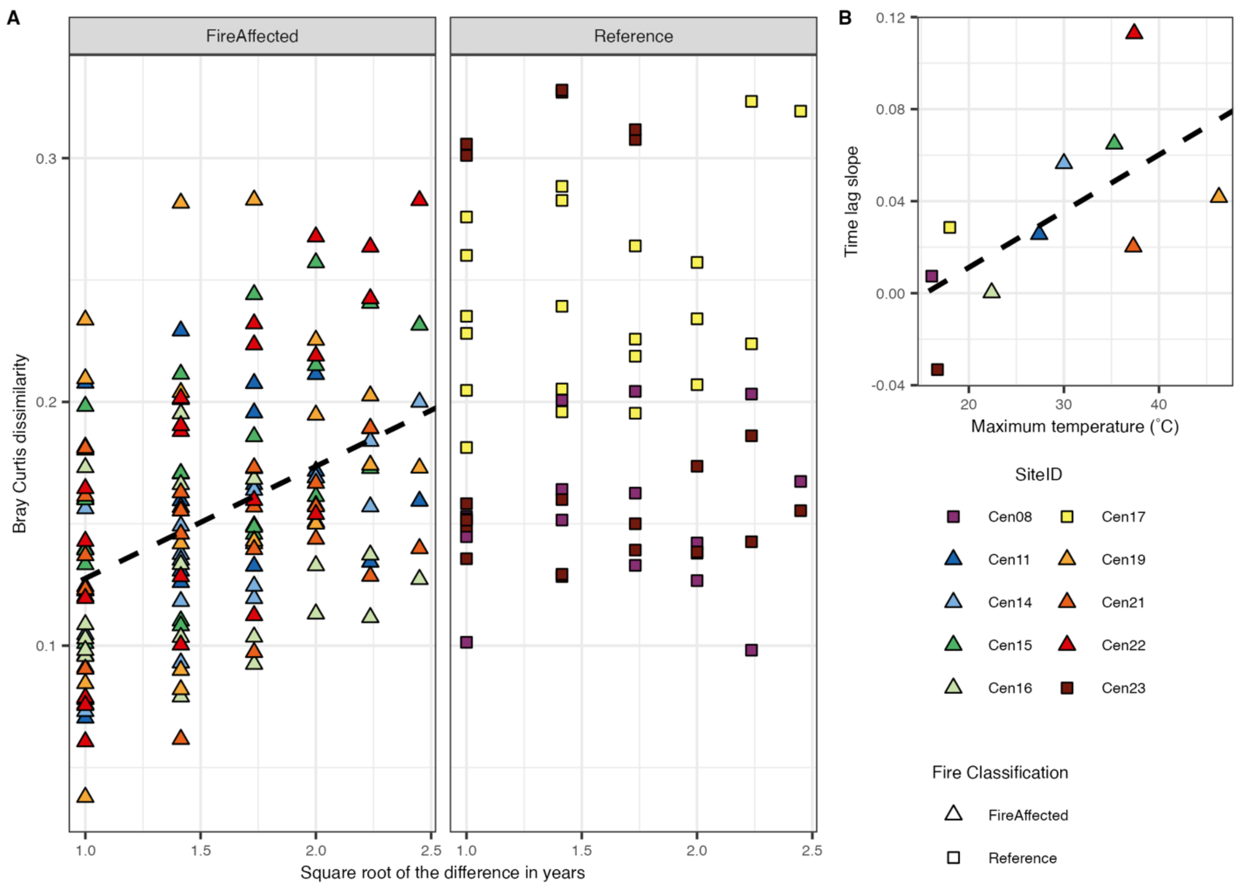


**Figure S5:** Timelag analysis measuring interannual variation in metagenome structure (KEGG orthologues). A) Variation in metagenome structure using Bray-Curtis dissimilarity between years within a site. The x-axis indicates the square root of the years between sampling timepoints (e.g., 2016 to 2018 = 2 years). Dashed line indicates the linear mixed effects regression applied to the data from each fire classification. B) The slopes of the time lag analysis for each site compared to the maximum soil temperature measured at the site (*i.e.,* disturbance intenstity). Dashed line indicates the linear regression applied to the data.

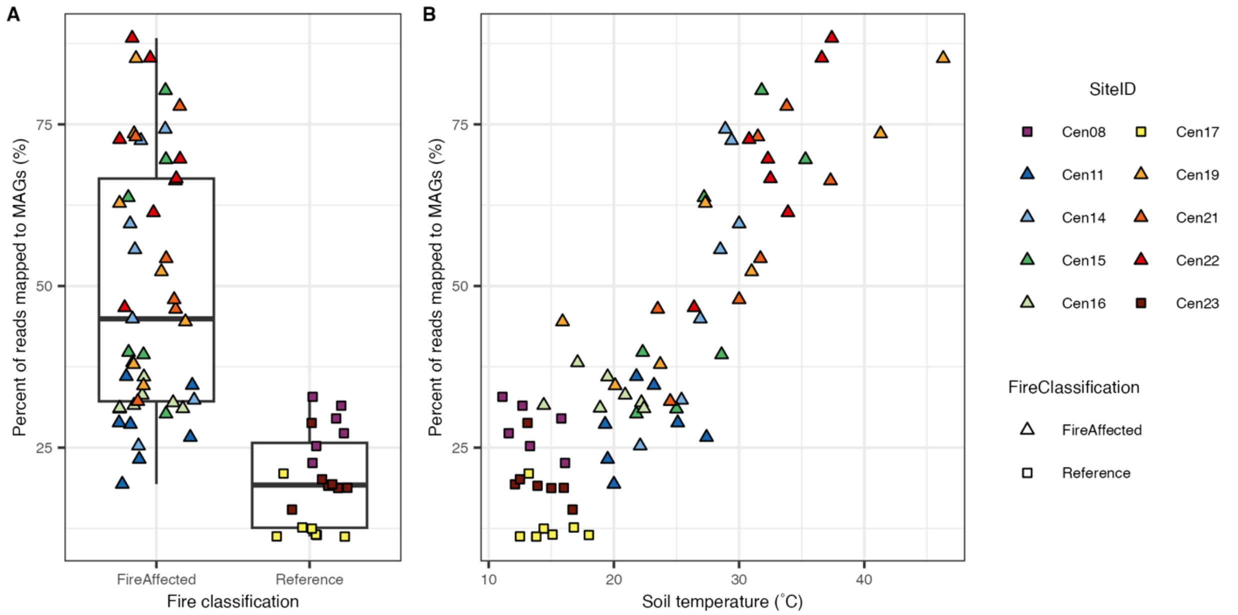


**Figure S6:** Percent of metagenome reads mapped to MAGs across A) fire classification and B) temperature.

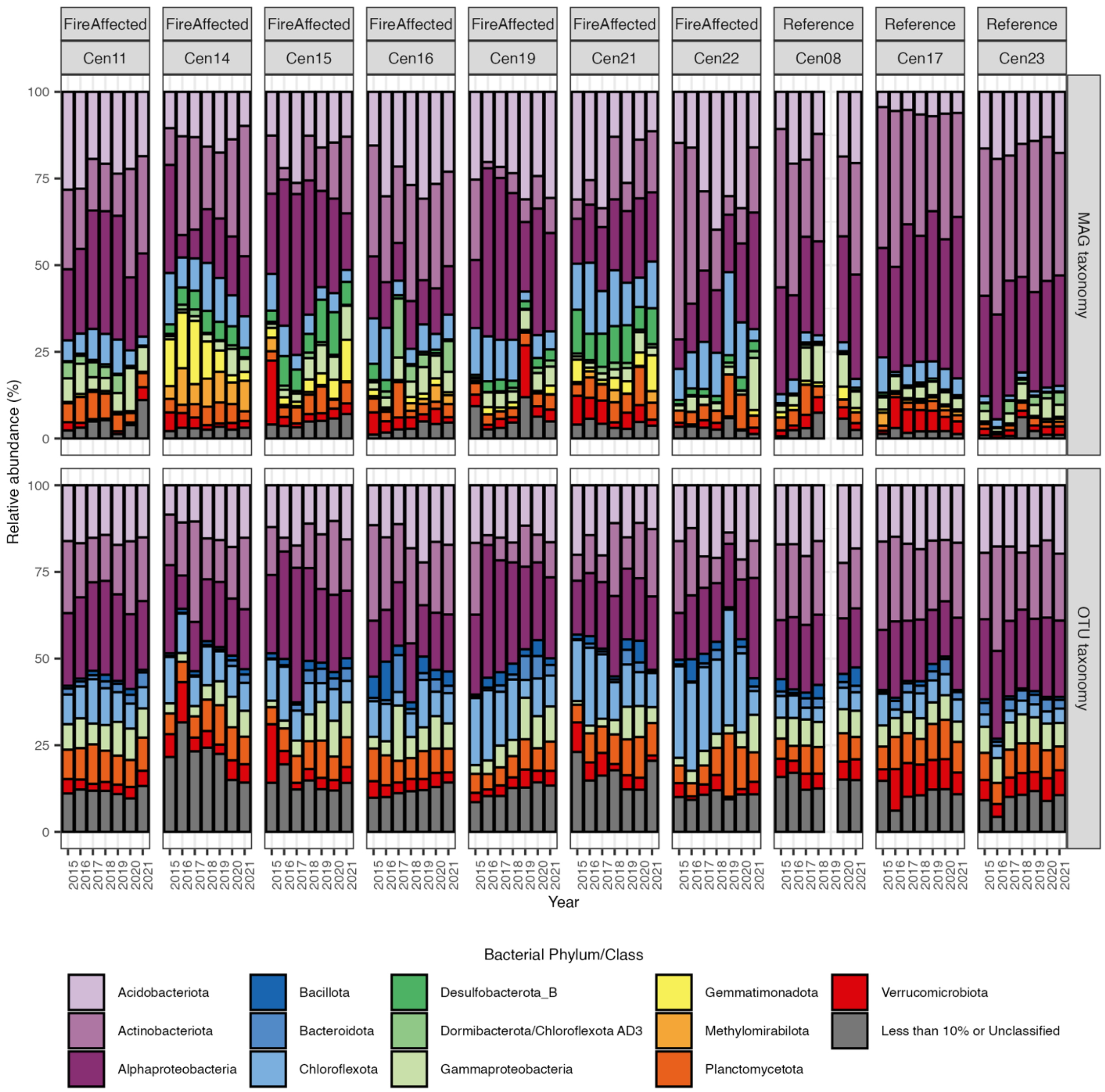


**Figure S7:** Comparison of the top most abundant MAG and OTU phyla (class for *Pseudomonadota*). Phyla that make up less than 10% of the dataset were combined with the taxa unclassified at the Phylum level.

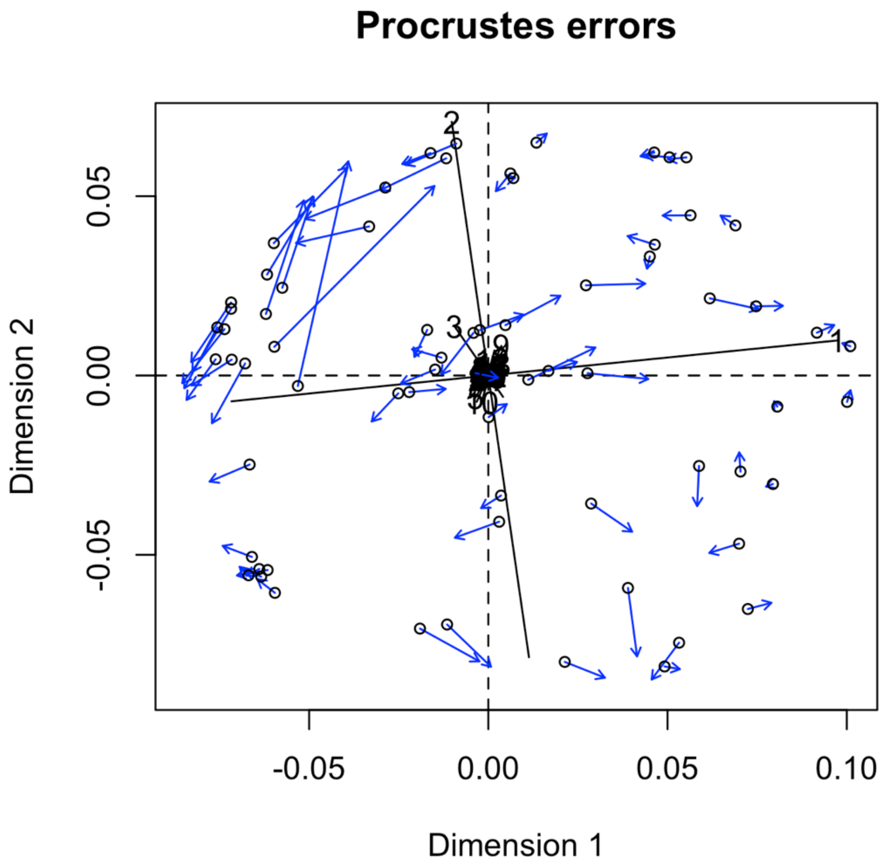


**Figure S8:** Plot of Procrustes errors between MAG based and OTU based community structures. Ordinations used were based on Bray-Curtis dissimilarity and PCoA.

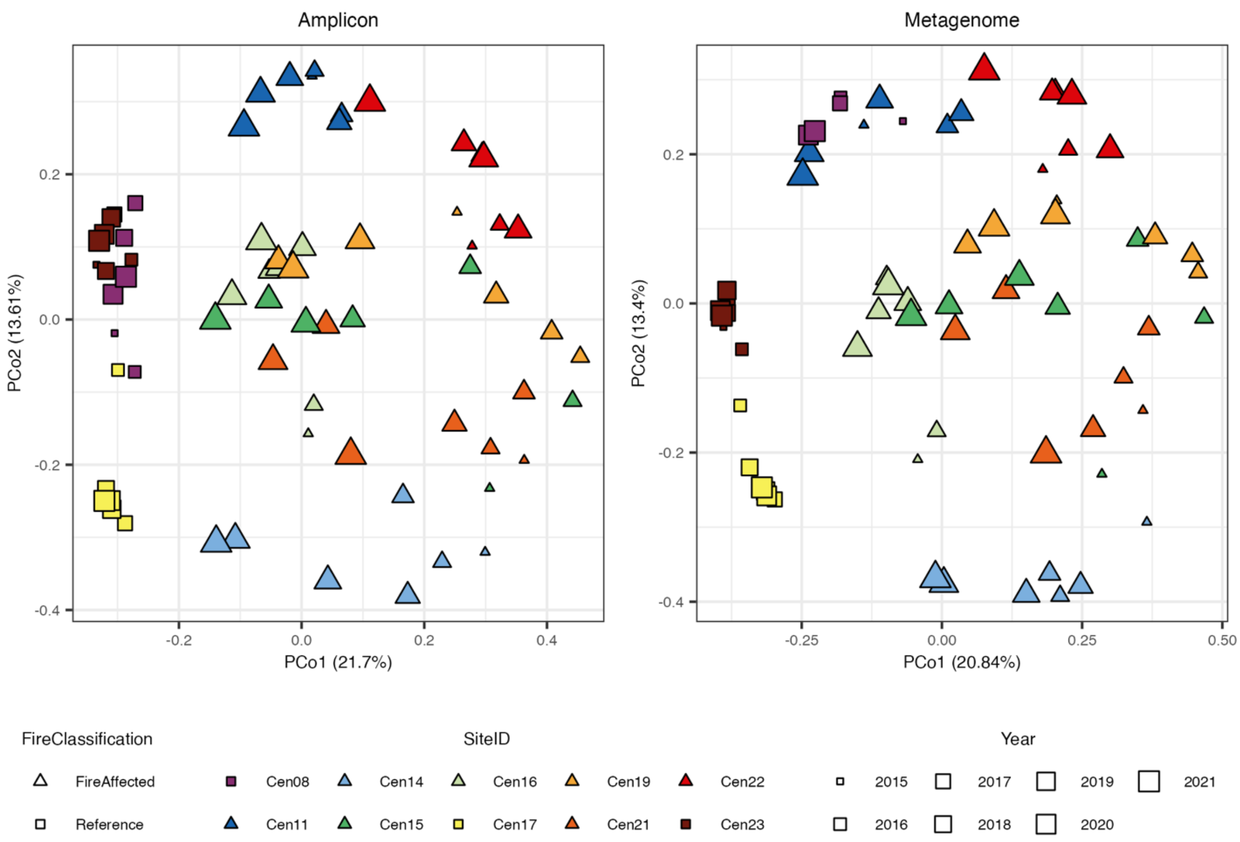


**Figure S9:** PCoA of Amplicon (OTU) and MAG based community structures. Both utilized Bray-Curtis dissimilarity.

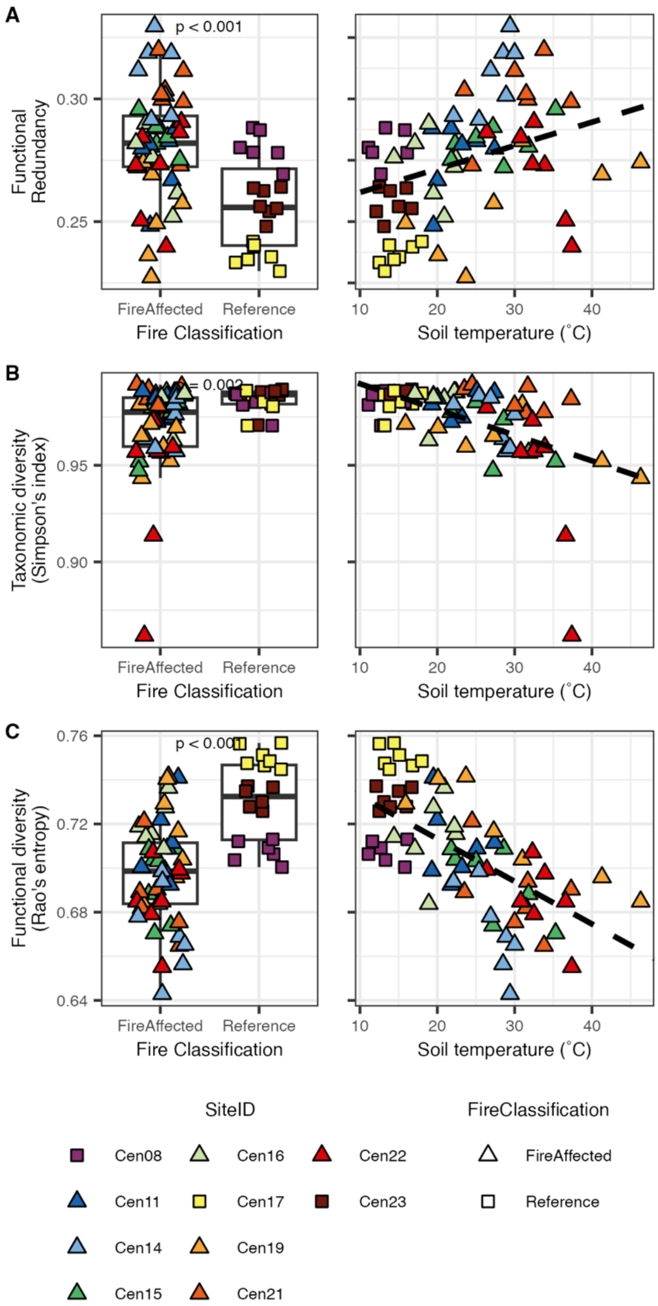


**Figure S10:** Functional redundancy as measured by the relationship between taxonomic diversity (Simpson's index) and functional diversity (Rao's entropy) using Gapseq pathways as functions. Results are similar to those based on KEGG orthologues (Fig. 2). A) Functional redundancy is higher in fire-affected soils than reference and decreases as soil temperature decreases. B) Taxonomic diversity is higher in reference than in fire-affected soils and increases as soils cool off. C) Functional diversity is higher in reference than in fire-affected soils and increases as soils cool off.

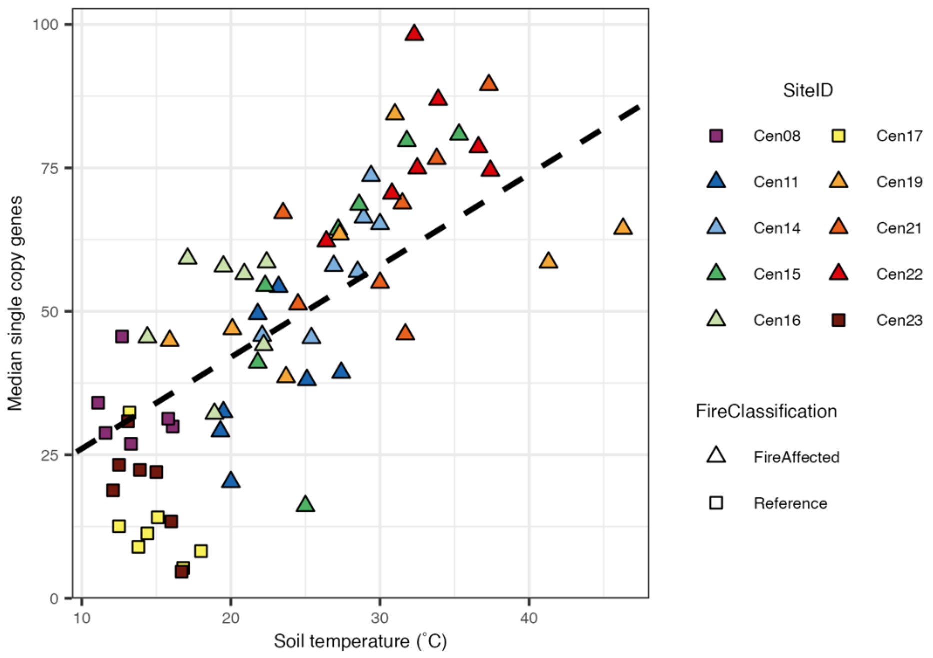


**Figure S11:** Median counts of universal single copy genes assembled and annotated in each sample compared to soil temperature. 35 ribosomal proteins and nine other universal single copy genes were examined from each assembly. Dashed line represents the linear mixed effects regression.

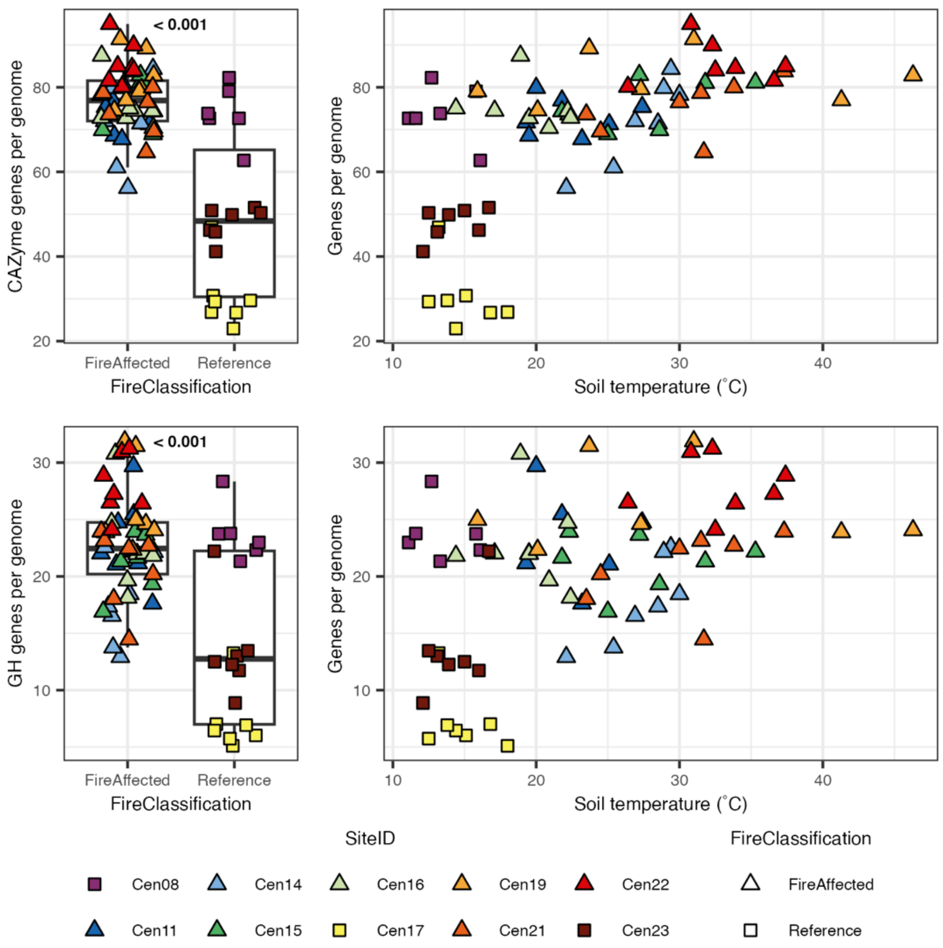


**Figure S12:** Per-genome investment in CAZyme and glycoside hydrolase (GH) genes. Pathway investment is calculated as the total number of open reading frames that mapped to CAZymes or GH from the dbCAN HMM database using HMMER divided by the median number of universal single copy genes also identified. For the boxplots, the p-value for the Wilcoxon test is indicated above the points.

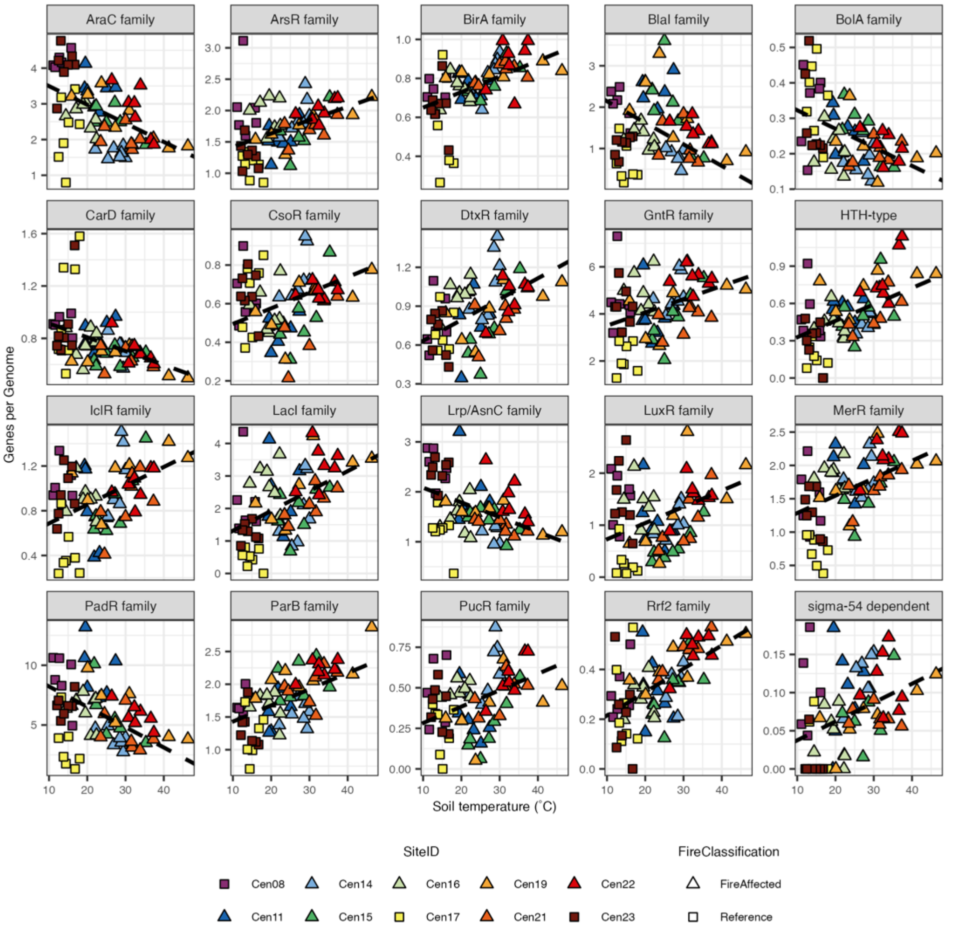


**Figure S13:** Per-genome investment in transcription factor families. Family investment is calculated as the total number of KEGG orthologues within each family detected across open reading frames within each sample divided by the median number of universal single copy genes also identified. Note that the y-axis indicating genes per genome varies in range by family.
